# DynaTMT: A user-friendly tool to process combined SILAC/TMT data

**DOI:** 10.1101/2021.07.06.451268

**Authors:** Kevin Klann, David Krause, Christian Münch

## Abstract

The measurement of protein dynamics by proteomics to study cell remodeling has seen increased attention over the last years. This development is largely driven by a number of technological advances in proteomics methods. Pulsed stable isotope labeling in cell culture (SILAC) combined with tandem mass tag (TMT) labeling has evolved as a gold standard for profiling protein synthesis and degradation. While the experimental setup is similar to typical proteomics experiments, the data analysis proves more difficult: After peptide identification through search engines, data extraction requires either custom scripted pipelines or tedious manual table manipulations to extract the TMT-labeled heavy and light peaks of interest. To overcome this limitation, which deters researchers from using protein dynamic proteomics, we developed a user-friendly, browser-based application that allows easy and reproducible data analysis without the need for scripting experience. In addition, we provide a python package that can be implemented in established data analysis pipelines. We anticipate that this tool will ease data analysis and spark further research aimed at monitoring protein translation and degradation by proteomics.

## Introduction

In the recent years, the usage of pulsed stable isotope labeling in cell culture (pSILAC) has grown to a state-of-the-art method to profile protein dynamics^1,2^. The combination with tandem mass tags (TMT) to multiplex samples further enhanced the throughput and sensitivity of these proteomics experiments^3–5^. Assessment of protein dynamics adds an important level of study to understand cellular responses to stimulation and has led to high impact discoveries in all areas of biological research, including basic cell biology and infection^5–8^. In those experiments, cells or animals are shifted towards feeding with isotopically labeled (i.e. heavy) SILAC amino acids for a defined period of time^9^. These heavy amino acids incorporate into newly synthesized proteins to make these distinguishable from the pre-existing “old” proteome that is not isotopically labeled (i.e. light). Quantification of the ratio of heavy versus light (that is “new” versus “old”) peptides then allows measurement of protein synthesis and degradation^1^. In addition, the combination of SILAC and TMT (SILAC/TMT), also called “hyperplexing”, has been introduced to greatly expand the multiplexing capacity of proteomics experiments^10^. SILAC/TMT experiments share the use of two introduced labels (a metabolic label [SILAC] and a chemical labeling reagent [TMT]) that are used to profile different conditions or protein species resulting in high data complexity that needs to be dissected after data acquisition. While the experimental design and workflow of SILAC/TMT experiments has become well established and available to a large number of researchers, data analysis remains tedious. After processing of the RAW files and peptide identification, the extraction and separation of the different labels from the TMT data requires either custom scripts implemented in analysis pipelines or tedious manual manipulation of data tables. This has limited the broad use of proteome dynamics experiments.

Here, we developed a browser based standalone tool called “DynaTMT” that enables rapid SILAC/TMT experiment data analysis. DynaTMT uses a user-friendly interface, compatible with Windows and MacOS environments, that returns readily processed data, including different normalization options, and basic visualization for quality control. For easy user access, DynaTMT does not require scripting experience or preinstalled tools other than a web browser. Our software is able to process multiplexed enhanced protein dynamics mass spectrometry (mePROD)^5^ data, implementing the previously published data analysis workflow^5,11^, or any other SILAC/TMT combination, including hyperplexing^10^, not requiring the additional calculation steps from mePROD. In addition to the user interface, we provide a python package to analyze SILAC/TMT datasets for integration into already established data analysis pipelines. We anticipate that these tools will open the field of protein dynamics to a wide range of researchers and extend its use towards new biological questions.

## Methods

### Implementations

#### DynaTMT browsertool

The DynaTMT browsertool was built using HTML, Jinja2, JavaScript and Python 3.8. Using the FLASK package, a python server starts processing and routing of the application. The server then uses the DynaTMT python package to process the input files. In addition to the DynaTMT package, the application uses the following packages: Python: Flask 1.1.2, Jinja2 2.11.2, numpy 1.19.4, pandas 1.1.4, pyinstaller 4.1, scipy 1.5.4, Werkzeug 1.0.1; JavaScript: Danfo 0.1.2, jquery 3.5.1, bootstrap 4.5.3, plotly.

The application in its distributed form does not need any pre-installed packages or programs, except a web browser like Firefox or Chrome. It is available from GitHub (https://github.com/klannk/DynaTMT) together with its source code and compiled versions for Windows and MacOS.

#### DynaTMT-py

The python package implementation of DynaTMT is available via GitHub (https://github.com/klannk/DynaTMT-py) and the PyPi repository (https://pypi.org/project/DynaTMT-py/) for python packages and is easily installable via the python package manager pip. It is intended to be implemented in already established pipelines and has some additional parameters that can be changed during analysis. The package is available under the GNU GPL v3 license.

### Injection time adjustment

To adjust TMT intensities for their injection time^11^, the TMT abundances are divided by the injection times by the following formula, where “i” is the TMT channel:

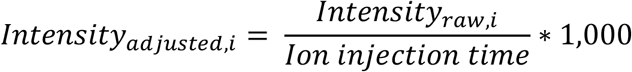

### Normalization

Three different modes can be used for normalization: 1) Total intensity normalization uses the summed intensity per TMT channel to calculate the normalization factors relative to the channel with the lowest total intensity. 2) Median normalization uses the median intensity of the channels instead of the sum. 3) TMM is a python implementation of the trimmed mean of M-values normalization introduced by Robinson et al.^12^.

### Baseline correction and protein quantification rollup

The baseline correction for mePROD experiments is performed as described before^5^: On peptide or perfect spectrum match (PSM) level (according to input), the abundance of the noise channel, which contains a sample that is not SILAC labeled, will be subtracted from all other samples^5^. This intensity is assumed to be generated from co-fragmented light peptides and can be considered as noise. To avoid artifacts generated by very small remaining intensities, we implemented a threshold value of 5 on average over all channels. After baseline correction, the mean of the channels has to be greater than the threshold value to be considered for further quantification. Otherwise, the peptide will be excluded. This threshold value can be fine-tuned empirically and set in the DynaTMT-py package. Negative intensities after correction will be set to zero to avoid negative numbers for noise measurements (the python package provides the possibility to change this behavior to either using random numbers between zero and one or keep the negative values). For the DynaTMT browsertool, protein rollup is performed by building the sum of all peptides/PSMs for a given unique protein identifier as this is the most implemented method in processing softwares like ProteomeDiscoverer (PD). In the python package, the method can be set to ‘sum’, ‘mean’ or ‘median’ to calculate protein quantifications. In addition the tool provides with peptide level tables as intermediate output to facilitate other downstream analysis.

### Re-analysis of available datasets

Primary processing of RAW mePROD data from our lab was carried out as originally described^5^.

PSM data was exported to a tab-delimited text file and further analyzed using DynaTMT. Hyperplexing data from Dephoure et. al. was kindly provided by the authors. We loaded the RAW files into PD 2.4 (ThermoFisher) and performed two SequestHT searches. Both searches included oxidation (M) as dynamic modifications and carbamidomethyl at cysteines as a static modification. One search node was used with fixed modifications for light peptides: TMT6 (N-term), TMT6 (K); and one search node for heavy counterparts: TMT6 (N-term), TMTK8 (+237.177, K). FDR was calculated using a target-decoy approach by Percolator. Reporter ion quantification was performed with TMT10plex settings. PSMs were exported to tab delimited text files and analyzed with DynaTMT.

## Results

### DynaTMT workflows

DynaTMT is compatible with any combination of SILAC and TMT or other isobaric labels, such as TMTpro or iTRAQ. Example setups are mePROD^5^ or hyperplexing^3,10^ experiments. Since mePROD workflows include an additional baseline channel, we included two preset workflows in the application (Figure 1). Both workflows are comprised of the same steps, except for the additional baseline subtraction during mePROD analyses and the optional ion injection time adjustment^11^. First, RAW files generated by the mass spectrometer are processed using the preferred software, such as PD. The default input files for DynaTMT are tables generated by PD. Nonetheless, DynaTMT accepts tab-delimited text files from any other processing software as well. However, when using other non-PD input files, the order of the columns needs to be adjusted for the correct information to be extracted by DynaTMT (see Supplementary Material 1). This behavior facilitates broad compatibility with any available processing software, such as MaxQuant^13^.

**Figure 1.**
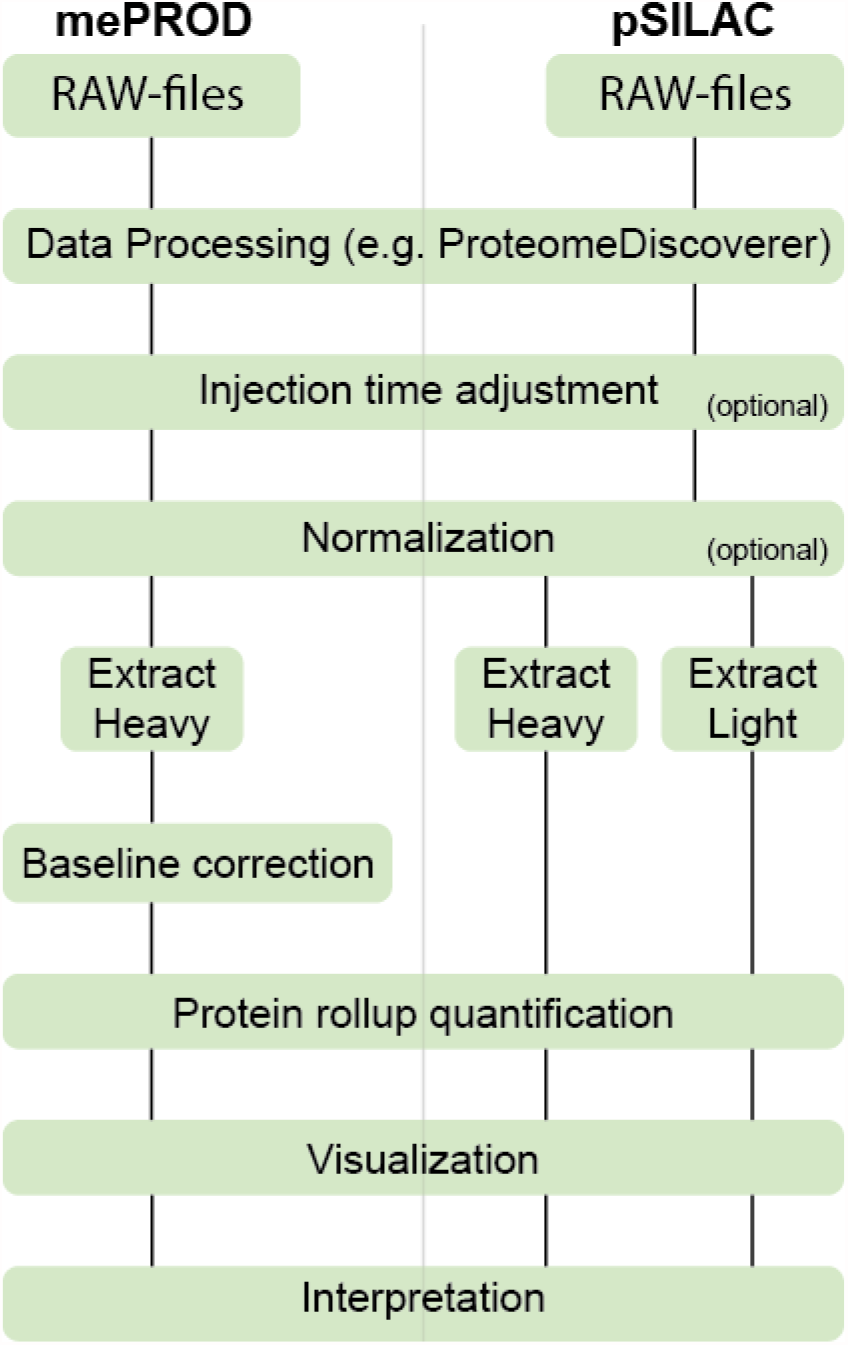
DynaTMT workflows. The application can be used for mePROD or any SILAC/TMT data, providing dedicated workflows for each approach. Both workflows are accessible from the application interface. After RAW file processing, the peptide data is read into DynaTMT and (optionally) adjusted for ion injection time. Subsequently, DynaTMT normalizes the data and extracts labels. Before protein rollup quantification during mePROD analyses, baseline substraction is required as an additional step. Data can be visualized using DynaTMT and exported as .csv files for further use.

The user can choose within the user interface (Figure 2) whether the data should be adjusted for the ion injection time. During SILAC/TMT experiments, ideally normalization happens using both heavy and light peptide TMT intensities. However, the TMT intensities do not necessarily scale with the peptide abundance. For high abundant peptides, the automatic gain control (AGC) target number of ions is filled rapidly while low abundant peptides need more time, but still might reach the same number of ions in the Orbitrap during the scan. Consequently, the different precursor abundances result in similar TMT quantification values. This becomes a critical issue during SILAC/TMT experiments since the workflow will create normalization artifacts overestimating heavy label abundance during translation measurements. Therefore, we implemented an ion injection time adjustment that has been shown to overcome this problem^11^. After injection time adjustment, the data is normalized using one of three normalization methods chosen from the user interface. We implemented three commonly used normalization techniques i) total intensity normalization, which is the default setting in most workflows, ii) median intensity normalization and iii) TMM normalization^12^.

**Figure 2.**
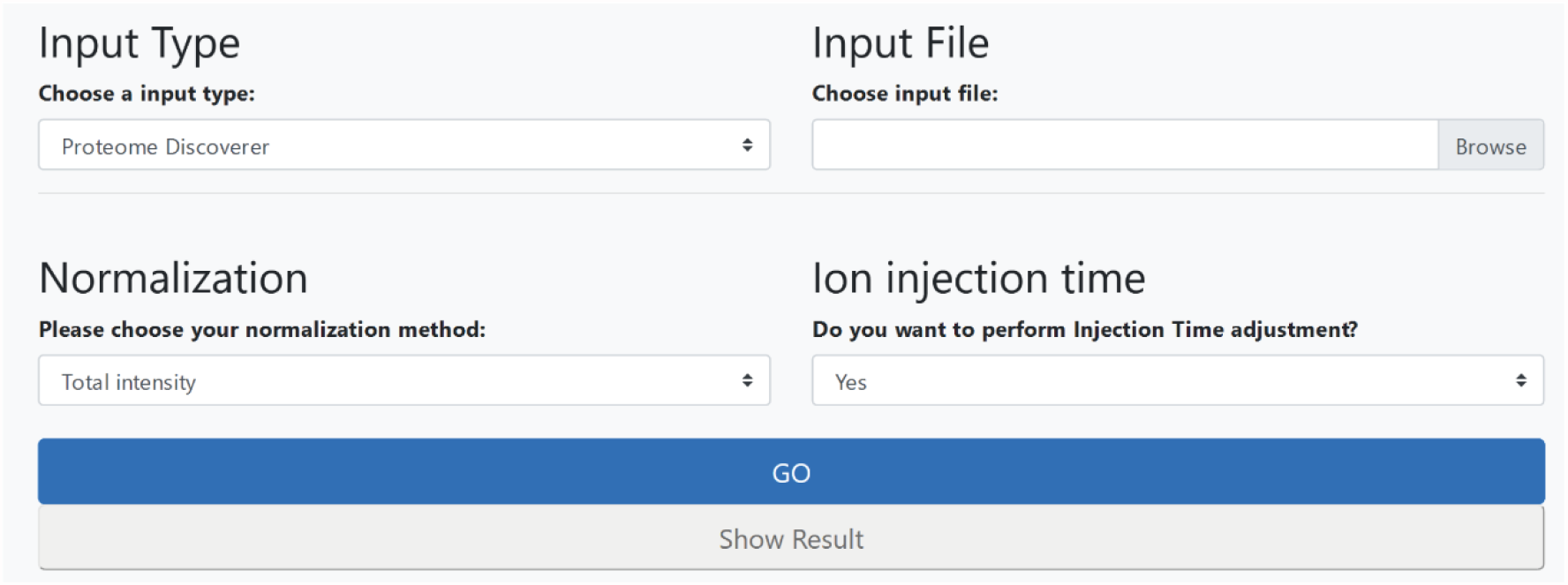
User interface of DynaTMT in the pSILAC mode. The user can choose the input file type (ProteomeDiscoverer, plain text file), the input file, one of three normalization methods (total intensity, median or TMM) and whether ion injection time adjustment should be carried out. The interface is identical to the mePROD interface, except for the lack to choose a baseline index channel, which is only applicable for mePROD data.

Normalization takes both heavy and light peptides in account assuming that the total protein amount inside a cell does not change during the experiment. After normalization, the peptides are sorted for their labels, extracted (heavy for mePROD, both labels for SILAC/TMT) into separate tables and quantification on protein level performed. While DynaTMT implements protein rollup via the summation of all peptides for a given protein, the python package has two additional options to calculate protein level quantifications: When calling the function either “sum”, “median” or “mean” can be selected to make the package more compatible with already established workflows.

For PD input files, DynaTMT automatically calculates statistical features for both heavy and light peptides individually, including average reporter ion intensity and isolation interference. This function is also available as a function in the python package, which generates a table with the relevant statistics.

### Reanalysis of available translation data

To evaluate the performance of DynaTMT, we reanalyzed a previously generated mePROD translation proteomics dataset using DynaTMT and compared to the original output^14^. To prepare the samples, HeLa cells had been treated with thapsigargin, an inducer of the ER unfolded protein response, or with a combination of thapsigargin and ISRIB, which reverses the effects of the UPR on translation. PSM data generated by PD was exported and analyzed using DynaTMT (Figure 3A). The analysis reproduced the global translation effects previously revealed^5^. As in the original analyses, we observed increased translation rates of known UPR-induced proteins, such as XBP1, HERPUD1 or HSPA5, further validating the capacity of DynaTMT to analyze mePROD data. Yet, differences in differential analysis expression were observed with DynaTMT when compared to the original publication. This is due to a change in the applied analysis workflow: The original mePROD publication did not apply a minimum threshold value that has to be reached after baseline subtraction. This resulted in a large number of proteins with a quantification value of zero or near zero. To facilitate better statistical downstream analysis, we implemented a threshold of average reporter intensity that has to be passed for each peptide to be included. However, this implementation leads to slight differences in the calculated fold change values comparing data analyzed with or without threshold. When we set the threshold to a negative value to include all quantifications, we were able to reproduce the fold changes from the original publication (Figure 3B). Notably, we also reproduced quantifications from data that was acquired using the targeted mass difference method and ion injection time adjustments (Figure 3C)^11^.

**Figure 3:**
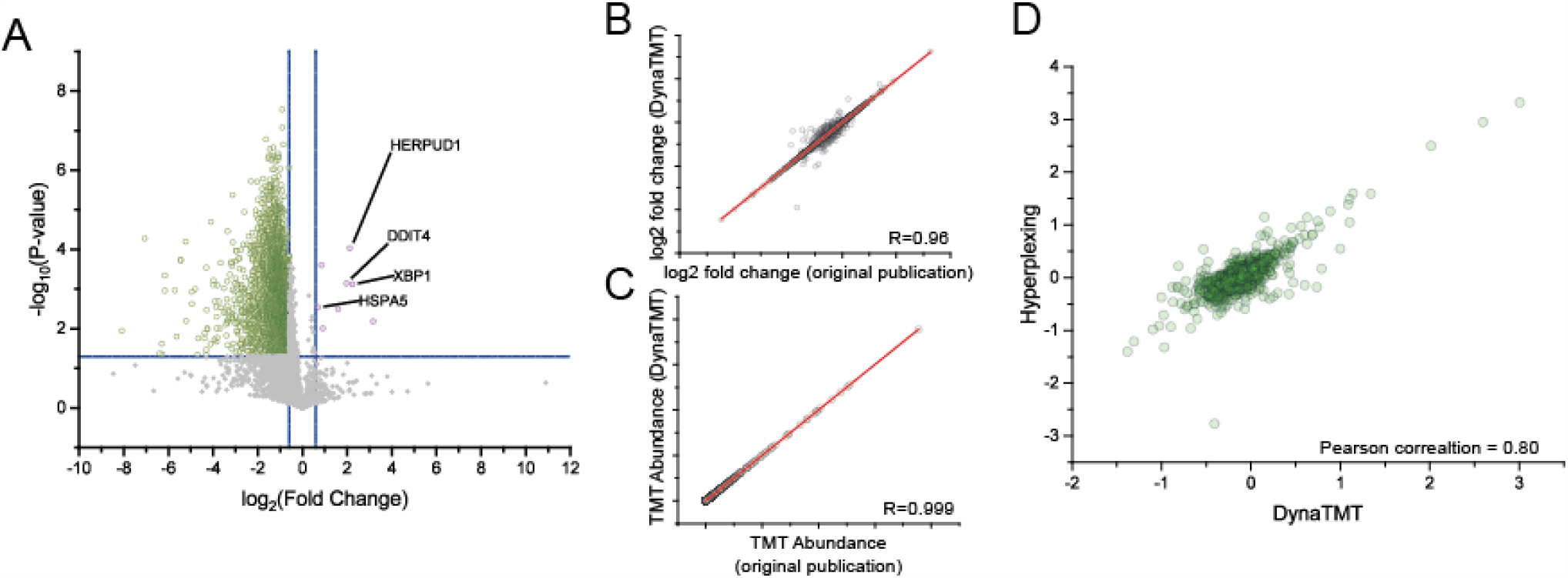
Reanalysis of available SILAC/TMT datasets for validation of DynaTMT. (A) Reanalysis of mePROD data. In the original publication, cells were treated with thapsigargin or DMSO and analyzed for changes in translation, measured by incorporation of SILAC amino acids. The reanalyzed data (DynaTMT browsertool) shows the expected induction of unfolded protein response genes, while global translation goes down. (B) Comparison of fold changes from the DynaTMT python package and the original publication. The python package was used to resemble the original data analysis workflow with the provided analysis options, since the original publication did not apply a threshold and used median based protein rollup. (C) Comparison of extracted TMT abundances from data using targeted mass difference based acquisition, together with ion injection time adjustments. (D) Fold changes from DynaTMT reanalysis and the original publication applying hyperplexing to increase TMT multiplexing capacity.

To demonstrate that DynaTMT is also compatible with any other SILAC/TMT hyperplexing data and to further evaluate its performance, we re-analyzed a dataset from Dephoure et al.^10^. In this dataset, the combination of SILAC and TMT was used to expand the multiplexing capacity of the TMT6plex reagents. In the reanalyzed dataset, yeast was subjected to rapamycin treatment and compared to control treatment with six replicates each. The addition of two SILAC labels in combination with TMT6plex allowed doubling the TMT multiplexing capacity to twelve. We reanalyzed the RAW files with PD 2.4. The peptide files were then loaded into DynaTMT and analyzed using the pSILAC mode with total intensity normalization and without ion injection time adjustment to better reflect the original analysis. Our obtained results correlated well with the original dataset (Figure 3D) with minor discrepancies likely driven by differences in downstream calculations of fold changes.

## Conclusion

The use of SILAC/TMT combination experiments to measure changes in protein synthesis and degradation is rising. While the processing of RAW files is nowadays performed by user-friendly software packages (e.g. ProteomeDiscoverer, MaxQuant, Spectronaut), the subsequent extraction of the SILAC data from the output files and the further processing (e.g. injection time adjustment) still requires custom scripting pipelines or manual table manipulations. To overcome this limitation and make SILAC/TMT experiments more accessible for researchers, we have developed DynaTMT. This application provides with the tools needed to perform reproducible mePROD and hyperplexing experiment data analysis within only a few minutes. Notably, a comparable tool for analysis of hyperplexing data was recently published^15^. While both tools have the analysis of hyperplexing data in common, our tools provides with the additional mePROD analysis workflow and importantly with an intuitive graphical user interface. Moreover, DynaTMT does not require additional quantification tools, except the ones from the initial processing softwares (e.g. PD or MaxQuant).

The availability of DynaTMT as both interface and python versions ensures its easy use and allows the straightforward analysis of protein dynamics profiling experiments. We anticipate that this tool will be a helpful resource to the field of proteomics and its applicators to increase the overall number of protein dynamics proteomics experiments carried out.

## ASSOCIATED CONTENT

### Supporting Information

Supporting information 1. Detailed documentation of the DynaTMT browsertool (PDF).

Supporting information 2. Detailed API documentation of the DynaTMT-py python package (PDF).

## AUTHOR INFORMATION

### Author Contributions

K.K. wrote the python script. The DynaTMT browsertool was written by K.K. with support from D.K.. The manuscript was written by K.K. with support from C.M.. C.M. conceived and supervised the study. All authors have given approval to the final version of the manuscript.

## ACKNOWLEDGMENT

We would like to thank the quantitative proteomics unit at Institute of Biochemistry II in Frankfurt. We also would like to thank the Noah Dephoure for kindly providing the RAW data. We further thank all testers of the software in the institute.

## ABBREVIATIONS

TMT: Tandem Mass Tag;
mePROD(-MS): multiplexed enhanced protein dynamics (mass spectrometry);
SILAC: stable isotope labeling in cell culture;
AGC: automatic gain control;
PSM: perfect spectrum match;

---

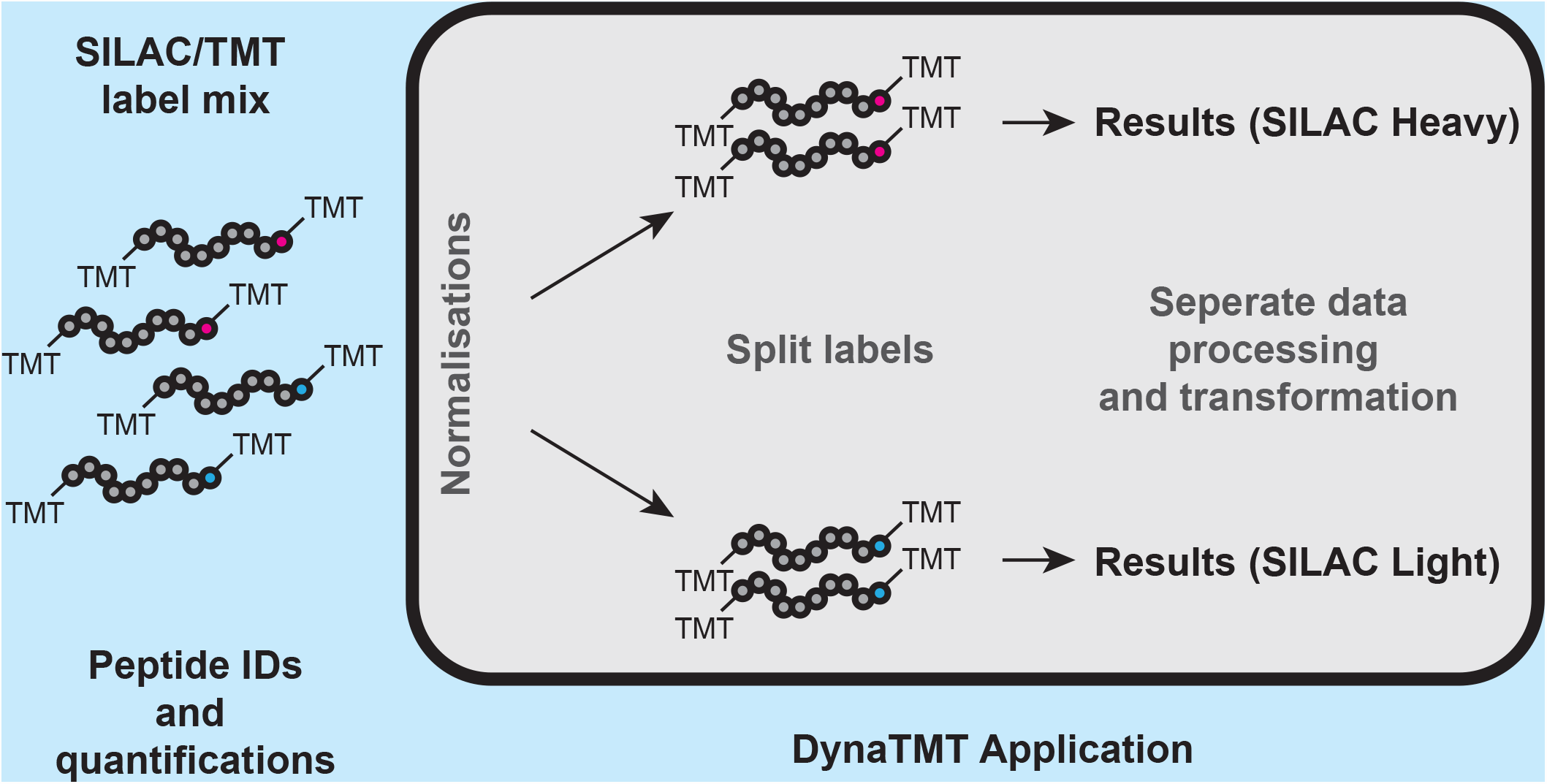

