## Supporting Information 2 for "DynaTMT: A user-friendly tool to process combined SILAC/TMT data"

### DynaTMT-py

The **DynaTMT tool** can be used to analyze **m**ultiplexed **e**nhanced **pro**tein **d**ynamic mass spectrometry (**mePROD**) data. mePROD uses pulse SILAC combined with Tandem Mass Tag (TMT) labelling to profile newly synthesized proteins. Through a booster channel, that contains a fully heavy labelled digest, the identification rate of labelled peptides is greatly enhanced, compared to other pSILAC experiments. Through the multiplexing capacity of TMT reagents it is possible during the workflow to use the boost signal as a carrier that improves survey scan intensities, but does not interfere with quantification of the pulsed samples. This workflow makes labelling times of minutes (down to 15min in the original publication) possible. Additionally, mePROD utilizes a baseline channel, comprised of a non-SILAC labelled digest that serves as a proxy for isolation interference and greatly improves quantification dynamic range. Quantification values of a heavy labelled peptide in that baseline channel are derived from co-fragmented heavy peptides and will be subtracted from the other quantifications. For more information on mePROD, please refer to the original publication Klann et al. 2020. The package can also be used to analyse any pSILAC/TMT dataset.

#### Install

```
pip install DynaTMT-py
```

#### Usage

##### Loading data

DynaTMT by default uses ProteomeDiscoverer Peptide or PSM file outputs in tab-delimited text. Relevant column headers are automatically extracted from the input file and processed accordingly. **Important Note:** DynaTMT assumes heavy labelled modifications to be named according to ProteomeDiscoverer or the custom TMT/SILAC lysinemodification, respectively. The custom TMT/Lysine modification isnecessary, since search engines are not compatible with twomodifications on the same residue at the same time. Thus the heavy lysine as used during SILAC collides with the TMT modification at the lysine. To overcome this problem it is necessary to create a new chemical modification combining the two modification masses. Please name these modification as follows:

- Label:13C(6)15N(4) – Heavy Arginine (PD default modification, DynaTMTsearches for Label string in modifications)
- TMTK8 – (Modification at lysines, +237.177 average modification mass)
- TMTproK8 - (Modification at lysines, +312.255 average modificationmass)

Alternatively, it is possible to input any other text file, derived from MaxQuant or similar programs, containing Protein Accession or Identifiers in the first column, Ion Injection times in the second column (optional) and Peptide/PSM Modifications in the third column (or second if no IT adjustment is performed). All following columns are assumed to be TMT intensities, no matter the column names. For these text files naming of the columns is irrelevant, as long as no duplicate column names are used. Alterantively you can change the all default column and modification names in the source code of the package if needed.

```
import pandas as pd

df = pd.read_csv("PATH",sep='\t',header=0)
```

### Workflow

mePROD uses injection time adjustment Klann et al. 2020 as a first step, but that is optional.

In the default workflow the Input class is initialised with the input data. This data is stored in the class and gets modfified during normalisation and adjustments.

#### IT adjustment

```
from DynaTMT.DynaTMT import PD_input,plain_text_input
processor = PD_input(df)
processor.IT_adjustment()
```

#### Normalisation

Normalisation is performed either by total intensity normalisation, median normalisation or TMM. Example (total intensity):

```
processor.total_intensity_normalisation()
```

#### Extraction of heavy peptides

Here a dataframe is returned by the function

```
extracted_heavy = processor.extract_heavy()
```

**If you use normal pSILAC TMT data without mePROD baseline channels, you can stop here and extract also the light data, by calling extract_light()**

#### Baseline normalisation

Here the baseline is subtracted from all samples and a dataframe on peptide level is created.

```
output =
processor.baseline_correction_peptide_return(input_file,threshold=5,i_baseline=0,r
andom=None)
```

##### Protein rollup

To create a protein level dataset, protein rollup will be performed by using one of the three implemented methods: 'sum', 'mean' or 'median'. Default is sum.

```
output = processor.protein_rollup(output,method='sum')
```

##### Store output

```
output.to_csv("PATH")
```

### API documentation

```
class PD_input
```

Class containing functions to analyze the default peptide/PSM output from ProteomeDiscoverer. All column names are assumed to be default PD output. If your column names do not match these assumed strings, you can modify them or use the plain_text_input class, that uses column order instead of names. **init**(self, input): Initialises PD_input class with specified input file. The inpufile gets stored in the class variable self.input_file.

```
IT_adjustment(self):
```

This function adjusts the input DataFrame stored in the class variabl self.input_file for Ion injection times. Abundance channels should contain "Abundance:" string an injection time uses "Ion Inject Time" as used by ProteomeDiscoverer default output. For other column headers please refer t plain_text_input class.

```
extract_heavy (self):
```

This function takes the class variable self.input_file dataframe and extracts all heavy labelled peptides. Naming of the Modifications: Arg10: should contain Label, TMTK8, TMTproK8 Strings for modifications can be edited below for customisation writes filtered self.input_file back to class

```
extract_light (self):
```

This function takes the class variable self.input_file dataframe and extracts all light labelled peptides. Naming of the Modifications: Arg10: should contain Label, TMTK8, TMTproK8 Strings for modifications can be edited below for customisation Returns light peptide Dataframe

```
baseline_correction(self,input,threshold=5,i_baseline=0,method='sum')
```

DECREPATED. Please use baseline_correction_peptide_return() instead and perform rollup with protein_rollup() function. This function takes the self.input_file DataFrame and substracts the baseline/noise channel from all other samples. The index of the baseline column is defaulted to 0. Set i_baseline=X to change baseline column. Threshold: After baseline substraction the remaining average signal has to be above threshold to be included. Parameter is set with threshold=X. This prevents very low remaining signal peptides to producartificially high fold changes. Has to be determined empirically.
Method: The method parameter sets the method for protein wolluquantification. Default is 'sum', which will sum all peptides for the corresponding protein. Alternatives are 'median' or 'mean'. Ifno or invalid input is given it uses 'sum'. Modifies self.input_file variable and returns a pandas df.

```
baseline_correction_peptide_return(self,input_file,threshold=5,i_baseline=0,random
=False,include_negatives=False)
```

This function takes the self.input_file DataFrame and substracts the baseline/noise channel from all other samples and returns a peptide level DataFrame. The index of the baseline column is defaulted to 0. Set i_baseline=X to change baseline column. Threshold: After baseline substraction the remaining average signal has to be above threshold to be included. Parameter is set with threshold=X. This prevents very low remaining signal peptides to producartificially high fold changes. Has to be determined empirically. By default negative values after baseline subtraction are replaced with zeros. For usage with linear models to avoid zero inflation two options exis: Either use include negatives = True, to avoid the replacement with zero values or use include_negatives = False and random=True to replace the values

```
protein_rollup(self, input_file,method='sum')
```

This function performs protein level quantification rollup by either summing all peptide quantifications or building the mean/median. method can be 'sum','mean' or 'median'.

```
statistics(self, input)
```

This function provides summary statistics for quality control assessment from Proteome Discoverer Output.

```
TMM(self)
```

This function implements TMM normalisation (Robinson & Oshlack, 2010, Genome Biology). It modifies the self.input_file class variable.

```
chunks(self,l, n)
```

```

```

Yield successive n-sized chunks from l.

```
total_intensity_normalisation(self)
```

This function normalizes the self.input_file variable to the summed intensity of all TMT channels. It modifies the self.input_file to the updated DataFrame containing the normalized values.

```
Median_normalisation(self)
```

his function normalizes the self.input_file variable to the median of all individual TMT channels. It modifies the self.input_file to the updated DataFrame containing the normalized values.

```
sum_peptides_for_proteins(self,input)
```

This function takes a peptide/PSM level DataFrame stored in self.input_file and performs Protein quantification rollup based on the sum of all corresponding peptides.

Returns a Protein level DataFrame

```
log2(self)
```

Modifies self.input_file and log2 transforms all TMT intensities.

```
class plain_text_input:
```

This class contains functions to analyze pSILAC data based on a plain text input file. The column names can be freely chosen, as long as all column names are unique. The different column identities are extracted by the column order:

1. Accession
2. Injection time (optional, is set in class init)
3. Modification

- all following columns are assumed to contain TMT abundances

  **init**(self, input, it_adj=True)

Initialises class and extracts relevant columns.The different column identities are extracted by the column order:

- Accession
- Injection time (optional, set by it_adj parameter)
- Modification
- all following columns are assumed to contain TMT abundances

**All other functions are used as for PD_input class**
