## Supporting Information 1 for "DynaTMT: A user-friendly tool to process combined SILAC/TMT data"

### Documentation

The **DynaTMT tool** can be used to analyze **m**ultiplexed **e**nhanced **pro**tein **d**ynamic mass spectrometry (mePROD) data. mePROD uses pulse SILAC combined with Tandem Mass Tag (TMT) labelling to profile newly synthesized proteins. Through a booster channel, that contains a fully heavy labelled digest, the identification rate of labelled peptides is greatly enhanced, compared to other pSILAC experiments. Through the multiplexing capacity of TMT reagents it is possible during the workflow to use the boost signal as a carrier that improves survey scan intensities, but does not interfere with quantification of the pulsed samples. This workflow makes labelling times of minutes (down to 15min in the original publication) possible. Additionally, mePROD utilizes a baseline channel, comprised of a non-SILAC labelled digest that serves as a proxy for isolation interference and greatly improves quantification dynamic range. Quantification values of a heavy labelled peptide in that baseline channel are derived from co-fragmented heavy peptides and will be subtracted from the other quantifications. For more information on mePROD, please refer to the original publication Klann et al. 2020.

*It can also be used to analyze any combination experiment of SILAC with TMT, if used in the pSILAC mode (see Navigation Bar). The documentation holds true for both workflows, just the baseline related calcualtions missing in pSILAC.*

#### Input Files

DynaTMT by default uses ProteomeDiscoverer Peptide or PSM file outputs in tab-delimited text. Relevant column headers are automatically extracted from the input file and processed accordingly.
**Important Note:** DynaTMT assumes heavy labelled modifications to be named according to ProteomeDiscoverer or the custom TMT/SILAC lysine modification, respectively. The custom TMT/Lysine modification is necessary, since search engines are not compatible with two modifications on the same residue at the same time. Thus the heavy lysine as used during SILAC collides with the TMT modification at the lysine. To overcome this problem it is necessary to create a new chemical modification combining the two modification masses. Please name these modification as follows:

- Label:13C(6)15N(4) – Heavy Arginine (PD default modification, DynaTMT searches for Label string in modifications)
- TMTK8 – (Modification at lysines, +237.177 average modification mass)
- TMTproK8 - (Modification at lysines, +312.255 average modification mass)

Alternatively, it is possible to input a tab-delimited text file from other sources containing Protein Accession or Identifiers in the first column, ion injection times in the second column (optional) and Peptide/PSM Modifications in the third column. All following columns are assumed to be TMT intensities, no matter the column names. For these text files naming of the columns is irrelevant, as long as no duplicate column names are used.

#### Normalization

DynaTMT normalizes the samples for their loading, based on both light and heavy peptides. It is assumed that the total protein level does not change between the conditions. The sample loading can be normalized using three different methods:

- Total intensity – Sum of all intensities in a channel
- Median – Median intensity of the channels is used to calculate normalization factors
- TMM – Trimmed mean of M values normalization (publication)

### Ion injection time adjustment

During TMT experiments, the resulting TMT intensity does not directly reflect the precursor abundance. Lower abundant peptides are injected longer to reach the same number of ions (the AGC target) in the orbitrap. Thus, especially in pSILAC experiments the normalization by just summing TMT abundances creates a bias in the quantifications (for more information please refer to this publication). To account for these differences it is possible to adjust the TMT intensities by their ion injection time. If you want to adjust your data, please make sure to export the ion injection times for your PSM file.

### Baseline Index

This field allows you to specify a custom index for your used baseline channel. Per default it uses the first channel. Please note that the array starts at 0. This means your first channel has the index 0.

### Results

The results are stored as tab delimited text files in the Results folder of the app. They are named according to the date and the time when the analysis was run.

### Visualization

From the dropdown menu in the visualizations tab, all results stored in the results folder can be accessed and boxplots of extracted TMT abundances are drawn. In case of standard ProteomeDiscoverer input, additional statistics like charge state, average reporter ion intensity or isolation interference are extracted and plotted for both heavy and light peptides. The plots are interactive and show relevant statistics and can be easily saved directly from the interface.
